# Social experience dependent plasticity of mouse *song* selectivity without that of *song* components

**DOI:** 10.1101/2021.10.26.466011

**Authors:** Swapna Agarwalla, Sharba Bandyopadhyay

**Affiliations:** Information Processing Laboratory, Department of Electronics and Electrical Communication Engineering, IIT Kharagpur; Advanced Technology Development Centre, IIT Kharagpur

## Abstract

Syllable sequences in male mouse ultrasonic-vocalizations (USVs), “songs”, contain structure -quantified through predictability, like birdsong and aspects of speech. Apparent USV innateness and lack of learnability, discount mouse USVs for modelling speech-like social communication and its deficits. Informative contextual natural sequences (SN) were theoretically extracted and they were preferred by female mice. Primary auditory cortex (A1) supragranular neurons show differential selectivity to the same syllables in SN and random sequences (SR). Excitatory neurons (EXNs) in females showed increases in selectivity to whole SNs over SRs based on extent of social exposure with male, but syllable selectivity remained unchanged. Thus mouse A1 single neurons adaptively represent entire order of acoustic units without altering selectivity of individual units, fundamental to speech perception. Additionally, observed plasticity was replicated with silencing of somatostatin positive neurons, which had plastic effects opposite to EXNs, thus pointing out possible pathways involved in perception of sound sequences.

## Introduction

Parallels between birdsong and speech have been a matter of interest for a long time^1,2^ because of similarities in vocal learning^3,45^ and production^6^, and presence of song and order selective auditory circuits^7,8^ in songbirds. Rudimentary components of the same have been argued to exist in the mouse^9,10^. However, mouse ultrasonic vocalizations (USVs), while context specific^11,12,13^ and of communicative significance^14,15^, appear to be largely innate^10,16,17^. Thus the mouse, a mammal with powerful genetic tools and techniques available^18,19^ fails to fully serve as the agent for a model for speech like communication and to study autism spectrum and other disorders with deficits in social communication^16^. Speech and social communication relies on informative sequences of sounds and *not*, significantly less informative, single isolated acoustic units^20^. The first step towards the above points would be to understand if mouse communication contains informative structured sequences of sounds. Simultaneously, it is equally important to know if its auditory system is capable of selectivity to structures of behaviourally relevant sequences as a whole and modify their representation based on social experience.

Selectivity to structural features of natural sounds^21,22,23,24^ species specific vocalization encoding^25,26,27^, plasticity of representation of vocalizations^28,29,^, fear or reward driven representational changes have all been studied in the A1 of the mouse^30^ and multiple species^31,32,33^. Encoding of simultaneously occurring sounds by spectral integration, like harmonics or other frequencies^30,34^ and selectivity to disyllables in marmosets^35^ have been limited to single sound tokens over short duration. However, studies on coding of sound sequences as whole, in A1, are missing.

In this study we primarily ask two questions. First, is the mouse A1 capable of representing an entire sequence of behaviourally relevant sounds as a whole object, showing selectivity to sequences? Second, does representation of such sequences show experience dependent plasticity without altering relative representation of components of the sequences? We use a sequence of contexts for male mouse USV production to record context dependent USV syllable sequences. Theoretically we show the presence of context dependent structure in sequences and derive informative sequences, called natural sequences (SN). Naïve female mice showed higher preference to SN than designed control random order syllable sequences (SR). Component syllables of sequences were found to be coded in naïve female mouse A1 single neurons in a differential manner is SN and SR, which remained unchanged in experienced (exposed to male) female mice. However selectivity of single neurons to SN and SR, probed through electrophysiology and 2-photon imaging, showed higher preference to SN that increased with degree of exposure. Excitatory neurons (EXNs) showed increased selectivity to SN over SR with exposure, while somatostatin positive inhibitory neurons (SOM INNs) showed the opposite behaviour. On the other hand parvalbumin positive INNs (PV) neither preferred SN or SR before, nor did their selectivity change with exposure. The same plastic changes in relative representation of SN and SR and not of component syllables are replicated with inactivation of SOM INNs paired with sequence presentation during and beyond pairing. We suggest that the known SOM INN based disinhibition of EXNs mediated by long duration activation of vasoactive intestinal peptide positive INNs (VIP)^36^ during the exposure experience.

Thus we provide the first evidence of selectivity and experience driven plasticity of representation of behaviourally relevant structure of entire sequence of sounds as a whole and not the components, in mice, akin to that in the songbird ^37,78^. Our results open up the possibility of establishing mouse communication as a model to study context based speech like communications with a set of components with predictive ordering, dependent on context ^13,38,39,40^. While many studies have used mouse USVs for behavioural phenotyping for a variety of neurodevelopmental and communication disorders ^16,41^, with basic features of single USVs and bouts, our results immensely increase the potential of mouse models to study such disorders. The current work also suggests a relook at studies on mouse vocalization production which have primarily focussed on production of syllables and syllable sequences without considering production of particular orders in syllables sequences.

## Results

### Male mice produce context dependent syllable sequences preferred by female mice

We developed a protocol for male mouse vocalization production, in three different contexts, referred to as Alone (M), Separated (MSF) and Together (MF) (Fig. 1A). Lone naïve adult male mice (*n*=4, P56-P90), with no social exposure to a female were placed in a cage in a sound chamber (M, 5-10 minutes) and they did not emit any vocalizations (Fig. 1A, *top left*). Next a naïve female mouse (P56-P90) was introduced into the cage, separated from the male by a mesh (Fig. 1A, *middle row*) and vocalizations were produced by the male (MSF, 5-10 minutes). Finally the mesh was removed and the female and male were exposed to each other and vocalizations were recorded (MF, 5-10 minutes). We repeated the process with the same pair of mice for at least 5 days until the male mouse emitted vocalizations in the Alone condition (see Methods). Ultra-frequency (UF) vocalization recordings (20kHz - 120 kHz) made from the above stage onwards for the following 5-7 days, in 4 pairs of mice, were further analysed and grouped into the three contexts stated above.

**Figure 1.**
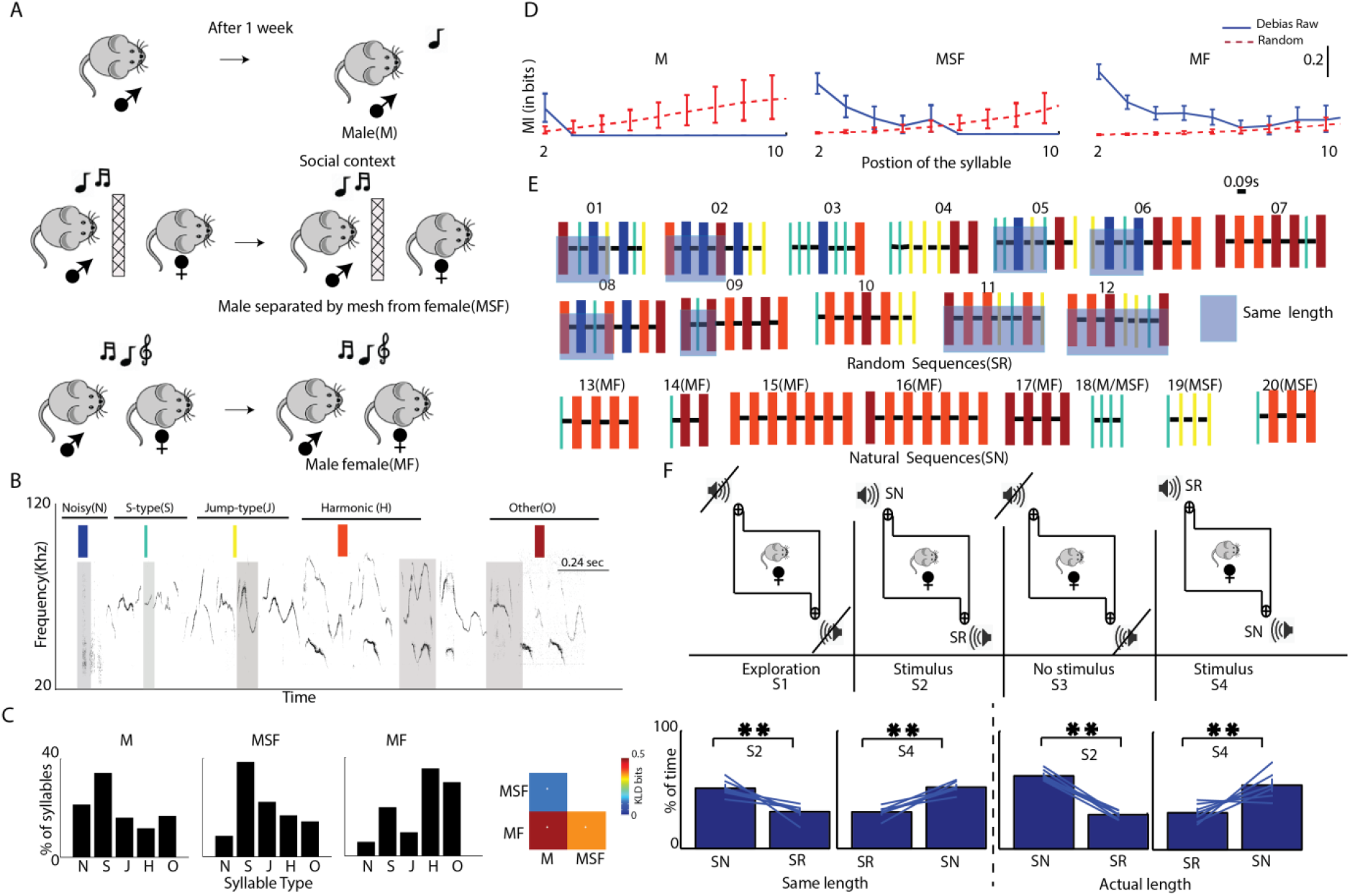
Naïve female mice prefer natural sequences emitted by male mice during social exposure. A) Schematic depicting the three social contexts (M, MSF and MF) in which male mouse vocalizations were recorded. B) Examples of spectrograms of representative syllable types, with one in each type highlighted with shaded background. Different syllable types are depicted with different color bars with widths denoting respective durations. C) Probability distributions, *pmfs*, of the different syllable types in the 3 contexts shown as bars with percentage of syllables. D) Plots of MI, *I*(*S*_*1*_; *S*_*i*_) with *i*=1,2 … 10 in the 3 different contexts in blue with 95% CIs. Red plots, with 95% CIs, show the expected extent of ‘0’ MI estimate from the data after scrambling order of syllables. Lack of overlap of the CIs (red and blue) indicate significant MI. E) The set of sequences created for SR and extracted for SN are depicted with the colored bars, as in B. The light blue background in a subset of the SR indicates the sequences for same length case. The SNs in 3 different contexts are identified above each SN. F) *Top row*: S1-S4 depicts the 4 sessions recorded with S1 and S3 having no sounds played. A fifth session S5, not depicted, was also recorded in which no stimulus was played. S2 and S4 have sounds played from speakers as indicated. *Bottom row*: The two bar plots to the left of the dashed line show bars indicating time spent on side of SN and SR in S2 and S4, in equal length case. The same for the SR having 7 syllables is shown to the right of the dashed line.

Vocalization sequences were first automatically annotated by detecting UF-syllables based on energy threshold (see Methods) and further by categorizing syllables into five types (see Methods, Fig. 1B, ^42^ based on their spectrograms with pitch jumps as the distinguishing feature^43^.The syllable types were, N (Noisy), S (Single frequency contour), J (Jump type, with a single jump in frequency), H (Harmonics) and O (Others – consisting of multiple Jumps). The relative percentage of the different kinds of syllables produced by male mice in each context (Fig. 1C) showed distinct differences. The overall difference in the distributions of syllables in each pair of the contexts was quantified using Kullback Leibler Divergence (KLD, see Methods)^44,45,42^. All pairs of distributions were significantly different at 95% CI (see Methods). However, the MF condition showed the most distinct distribution having high distances from the others, with the M and MSF cases showing a similar nature. In spite of differences in distributions, order of syllables could be random in the 3 contexts. To probe the non-random nature of syllables, we analysed syllable to syllable transition probability matrices as done in multiple studies^39,43,46^(see Methods) with two successive syllables (Fig. S1A) or disyllables. We found distinct differences, quantified with KLD (Fig. S1B), between the different contexts suggesting differences in structure of syllable order.

In order to probe higher order structure in syllable sequences, analyses as above with 3 or more successive syllables require large amounts of data for reliable estimation of transition probabilities. Thus to analyse structure in the order of syllables emitted^39,42,43,47^ the entire period of vocalization recordings of each session was divided into bouts based on the inter-syllable interval distribution (Methods, mean=90 ms, STD=160 ms). Silence intervals more than 250 ms in duration were used to mark the end of a bout of vocalizations (Fig. S1C). The syllable following the immediately preceding end of a bout was considered as the start of the next bout and successive syllables in a bout were considered as a vocalization sequence of syllables^42^. Thus bouts of vocalizations could have only one or many syllables (observed up to 30 syllables, Fig. S1D, total 948(M), 3320(MSF) and 6656(MF) bouts recorded). Bouts of vocalization with 3 or more syllables are referred as *sequences* (see Methods) in the rest of the paper. With *S*_*i*_ denoting the *i*^th^ syllable of a bout, the mutual information (MI) between *S*_*1*_ and *S*_*i*_, denoted *I*(*S*_*1*_;*S*_*i*_) were computed, debiased and tested for significance based on CI (see Methods). Significant MI between syllables at different positions in a bout shows dependence between the syllables or predictability, the basis of structure in sequences^20,42^. We found that in the 3 contexts significant dependence existed between the first and other syllables in bouts of vocalizations (Fig. 1D). The dependence was strongest and longest in the MF context and weakest and shortest in the M condition. Thus the 3 contexts had different degrees of dependence in syllables of sequences. Similar results were observed for *I*(*S*_*j*_;*S*_*j+k*_) for *j* = 2,3 … and *k* = 1,2 … (Fig. S1E) showing dependence exists within the sequence beyond that on the first syllable. Thus context dependent sequences are produced by male mice with degrees of structure varying with context.

To further understand what order of syllables, if any, were involved in providing such dependence in the sequences, we found the high probability sequences produced in the 3 contexts (Fig. 1E, sequence 13 to 20, see Methods). The 8 sequences obtained are called natural sequences, henceforth and denoted by SN. The natural sequences or their sub-sequences (from the starting syllable with at least 3 syllables) constituted 36% (MF) and 24% (MSF) of all the recorded sequences of length 3 or more in the two contexts. The percentages of such sequences occurring by chance based on the MF and MSF syllable probability distributions (Fig. 1C) were only 11% and 8%. The SN were also 8 of the 10 sequences with the highest accumulated surprise^48^ see Methods) showing that they were among the most informative^44^ among the repertoire of syllable sequences produced in the 3 contexts. As expected from the results of MI (Fig. 1D), the longest SN were in the MF condition and MF and MSF conditions had distinct sets of SN indicating context dependent specific sequence production in social encounters of male and female mice. The only high probability sequence in the M context was also present in the MSF context and is possibly a search sequence prior to finding a female mouse. However, all the SN sequences were present in the MF condition and thus during MF the female was exposed to all the SN. To study the importance of the SN, both from a behavioural perspective and coding perspective, as control, 12 other sequences were designed. Four randomly ordered sequences were created from the probability distributions of syllables in each context (Fig. 1C). The above sequences each with 7 syllables, the length of the longest SN, are considered as random sequences (Fig. 1E, sequence 1 to 12, see Methods) and denoted by SR.

Although SN were obtained from male vocalizations produced during social exposure to females, the sequences need not be behaviourally relevant to naïve females. We tested the relevance of the specific sequences obtained with a 2-sided free access/choice test^12,39^(see Methods). Two sets of experiments were done, one in which 8 of the 12 SR were truncated to match the number of syllables in SN (Fig. 1E, shaded SRs, same length) and the other in which all SR had 7 syllables as in the longest SN. The latter case was used to provide the maximum possibilities of syllable to syllable transitions that could be naturally occurring. The above allowed us to probe the relevance of the full sequence as opposed to natural transitions or disyllables occurring randomly. Naïve female mice (P56-P90, n= 9 (2 outliers), same length, n=7 (1 outlier), 7 length) were placed in an open cage, for 4 distinct sessions (S1-S4) with sounds presented in S2 and S4 (see Methods). Sequences from SR were played back from one corner and those from SN from the opposite corner, alternately every 5 seconds, in S2. SR and SN’s respective sides were switched in S4 to remove any side bias (Fig. 1F, *top row*). Female mice had significantly higher preference (*p*<0.001, both cases) for SN independent of which side SN was played back, both for equal length and length 7 sequences (Fig. 1E, *bottom row*, quantified by dividing the cage floor into 3 equal parts denoting SN, Neutral and SR, see Methods). Thus the extracted natural sequences are relevant to naïve female mice and are preferred over the designed SR.

### Single units in A1 code single syllables and disyllables differentially in SN and SR

To investigate coding of behaviourally relevant natural sequences of vocalizations, SN, relative to SR, we first performed extracellular single unit recordings from Layer 2/3 (200-350 um from the surface) of mouse A1 using multi-electrode arrays ^49,50^ (Fig. S2AB). Three groups of mice were used, passively listening awake naïve females (AwF group, 328 units from 6 animals, chronic recordings, over 10-14 days), anaesthetized naïve females (AF group, 266 units from 12 animals) and naïve males (AM group, 195 units from 8 animals). Typically 5 (between 4 and 6) repetitions of each stimulus of SR (1-12) and SN (13-20) were presented in pseudorandom order (Fig. 2A, *left*) and single unit spiking (Fig. 2A inset, red) with 500 ms baseline was recorded. The stimuli varied in length based on the component syllables, from 0.389s to 1.233s. PSTHs were constructed and responses locked to single tokens were observed throughout the sequences of SR and SN (Fig. 2A, *right*).

**Figure 2.**
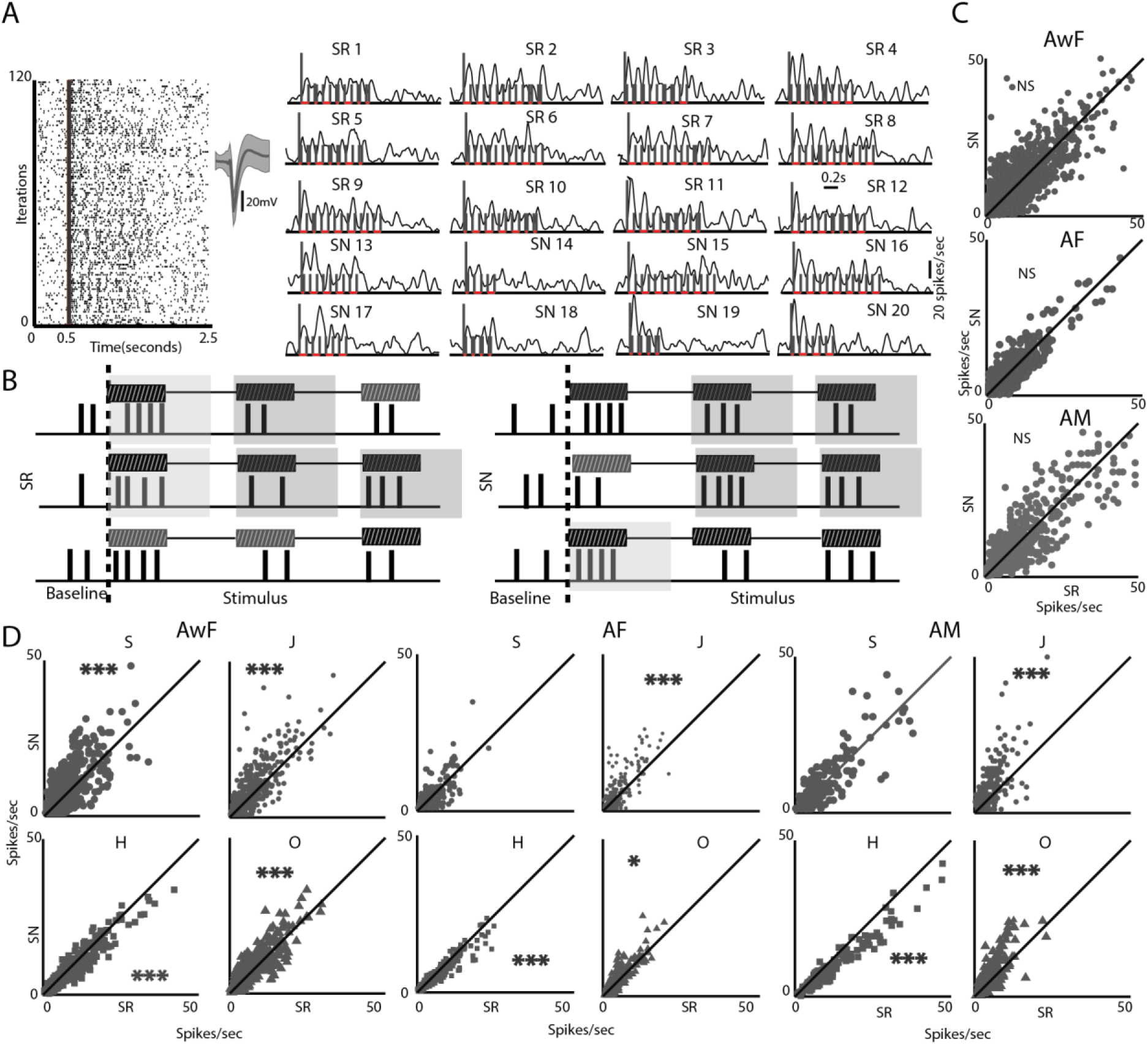
Coding of single syllables in mouse A1 depends on context. A) Representative dot raster plot of single unit spiking responses to SN and SR presented in pseudo random order (spike shape to the right). Smoothed PSTHs of the same unit for each stimulus sequence is shown with stimulus start (tall line) followed by lines marking start and end of each subsequent syllable. B) Schematic for calculation of responses to common syllables in the first position and within the sequences in SN and SR. C) Scatter plots show comparison of mean response rates of all common first syllables in SN and SR in 3 groups of mice, AwF, AF and AM. D) Scatter plots comparing mean response rates of common syllables (identified by solid symbols, S: large circle, J: small circle, H: square, O: triangle) in SN and SR, excluding occurrence in the first position for the same groups in C.

Syllables S, J, H and O and disyllables or transitions S-S, J-J, H-H and O-O were present in both SR and SN, excluding the first syllable any sequence. The first syllable (*S*_*1*_) was excluded as its occurrence does not contain any sequence information (Fig. 2B). Thus, as a control, in all 3 groups of animals average rate responses of each syllable type when occurring as *S*_*1*_ were compared (grey shade, Fig. 2B, see Methods). As expected, none of the syllable’s responses in the first position showed any difference between SN and SR (Fig. 2C, Fig. S2A for each syllable). Next, we compared the single unit rate responses of syllables and disyllables, present in both SR and SN, between the random and natural conditions (Fig. 2B, see Methods). Syllables J and O, across all three groups of animals, elicited higher average rates in A1 single units when occurring in SN than when they occurred in SR (*paired t-test*, *p*<0.01 in all cases, except syllable O in AF had *p*<0.05, Fig. 2D). However, syllable H across all the animal groups evoked lower average rates in SN than in SR (*paired t-test*, *p*<0.01 in all cases). Syllable S produced significantly higher rates when in SN (*paired t-test p*<0.01) compared to when in SR for AwF group and response strength of S was not significantly different in SN and SR in the other 2 groups of animals. Thus the single syllables were encoded differentially in a natural context than in a random context – specifically a higher response was generally elicited in SN than in SR, except for the H syllable. The differences could be an effect of stimulus specific adaptation SSA, 51,50,52 due to repeated presentations of syllables more so in SN (for example H in SN15 and SN16). Absence of such long repeats of H in SR could potentially explain the lower response rates to H in SN compared to SR (Fig. 2D). Repetitions of up to 4 successive syllables are present for S, J and O in SN with such repeated occurrence of them absent in SR, but these syllables evoked higher rates in SN than SR. Thus it is unlikely that adaptation can explain the observed context dependent responses to tokens. To finally rule out adaptation to be the cause of such differences, we compared the average adaptation profiles (Fig. S2B) in SN and SR, by normalizing each single unit’s mean response to the first token to be 1, which was not significantly different between SN and SR for any token (Fig. 2C, Fig. S2A). No significant differences were found in the adaptation profile based on average token wise responses over SN and SR across animal groups.

The above results suggest that relative ordering of the syllables is likely an important determinant of the difference in responses of individual syllables in SN and SR. We thus considered coding of the common disyllables in SN and SR, considering only the response to the second syllable in the transition (Fig. 3A, see Methods). Comparison of rate responses to transitions in SN and SR showed similar results as single syllables, with S-S, J-J and O-O transitions generally showing stronger responses when present in SN than when present in SR across all animal groups (except S-S in AF and AM, Fig. 3B–D). As with responses to the single syllable H, the transition H-H had higher responses in the random context compared to the natural context across all groups of animals (Fig. 3B–D). Syllable transitions of other kinds (Fig. S1A) were not common to both SN and SR. All the other kinds of disyllables in SN were present as the first transition, which were excluded, as the first syllable did not have context information (Fig. 2C and S2A). However, the other disyllables occurring as the first transition in SN (S-J, S-H, S-O and O-H) were significant in transition matrices (Fig. S1A). Thus we compared responses of the above transitions based on the response to the second token of the transition, between SN and SR (Fig. S3). In the above remaining cases also we generally find higher responses to the transitions when occurring in SN compared to that when occurring in SR.

**Figure 3.**
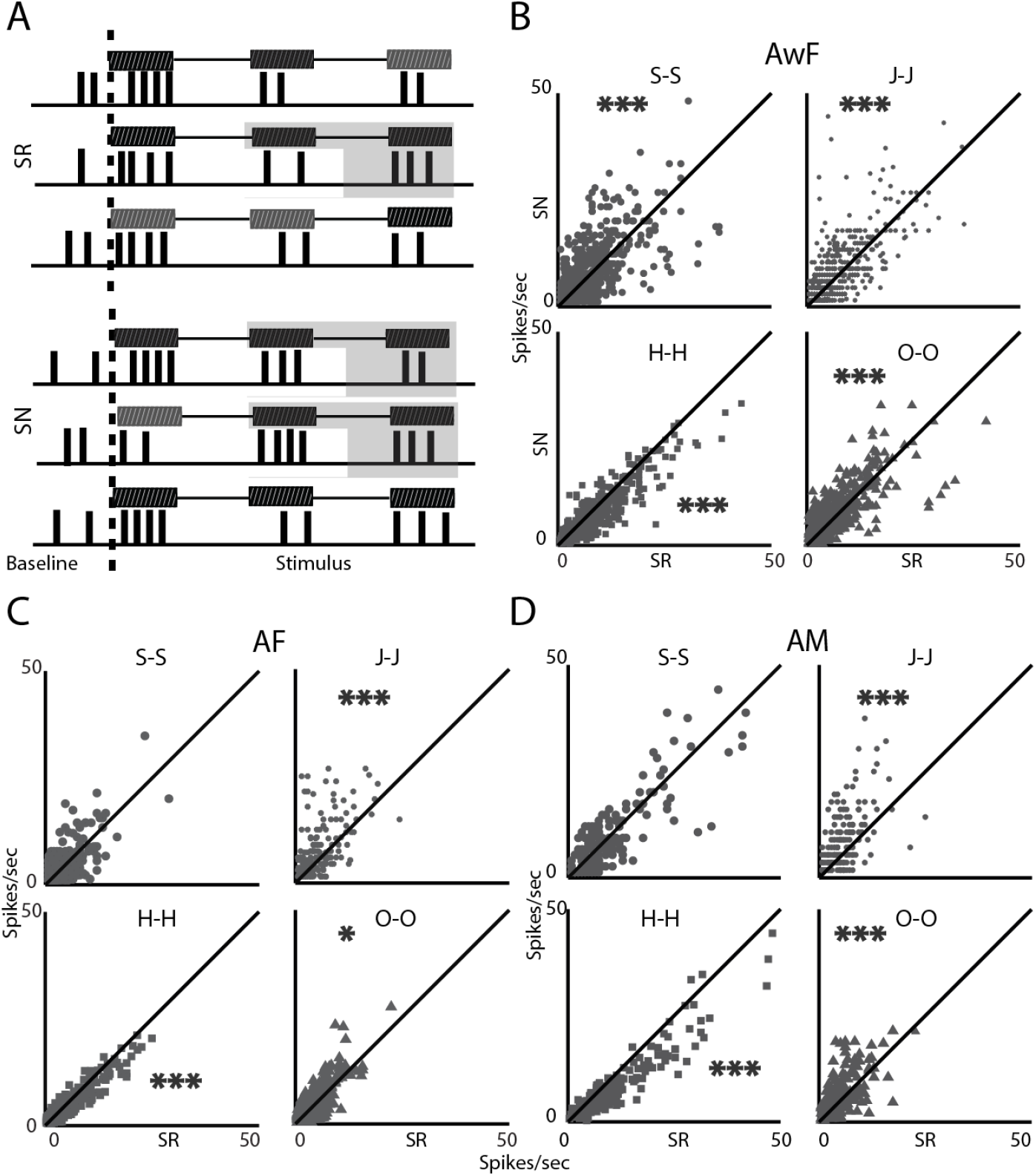
Coding of single disyllables in mouse A1 depends on context. A) Schematic for calculation of mean rate responses to transitions/disyllables based on response to the second component, excluding the first transition. B-D) Scatter plots comparing mean response rated of common disyllables in SN and when in SR for the three groups AwF (B), AF (C) and AM (D).

### Social exposure to male alters coding of entire natural sequences but not their component syllables in female A1

Our results show that naïve males and females both show the same results of differential coding of single syllables in the two sets of stimuli, SR and SN (Fig. 3). Further, the SN stimuli used are preferred by naïve female mice compared to SR (Fig. 1F). Social exposure between adult male and female mice (>P56) is a significant event constituting the SN USVs produced^15,39,53^ (Fig. 1). Further H syllables and sequence of H’s were a major component of the USVs especially in MF (Fig. 1C, 1E). Thus we hypothesize that the context specific coding of syllables and disyllables would change, especially for syllable H and disyllable H-H, both of which showed lower responses in SN.

We first tested the above with single unit recordings from Layer 2/3 of A1 of experienced female mice (Fig. 4A, see Methods) with different number of days of exposure to male mice (1-5 days of exposure, 421 units from 8 animals, AF-AFEx). Since the differential coding observed for syllables and disyllables were same in AwF and AF (Fig. 2C and Fig. 3BC) except for S, we considered rate responses in anaesthetized mice. First, as a control for context independence, we confirm that responses to all first syllables did not change between SN and SR (Fig. 4B, and Fig. S4A for each syllable type). Contrary to our hypothesis we found that experienced females had identical differential representation of single syllables (Fig. 4C) and disyllables (Fig. 4D) as the naïve females (Fig. 2C and 3B-D, before exposure, AwF-BFEx and AF-BFEx, except S and S-S in the latter case). However, selectively stronger responses to S in SN in AwF-BFEx and AF-AFEx compared to that in SR, suggests that the exposure likely had little effect on context specific coding of single syllables and transitions. As in the other groups, adaptation could not explain the context dependent coding of single syllables in the experienced females (Fig. S4B).

**Figure 4.**
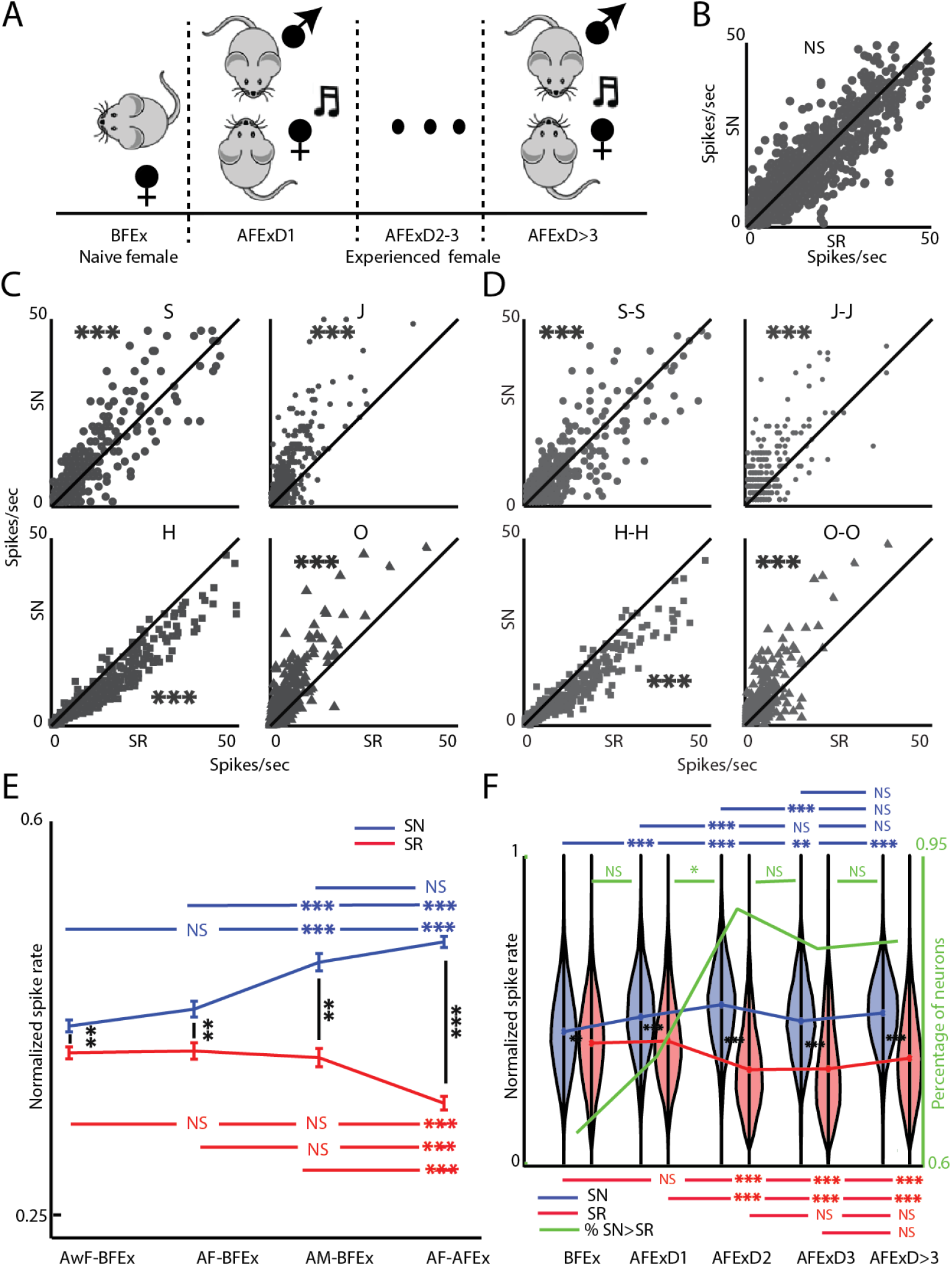
Plasticity in single units’ selectivity to entire SN sequences and not its components. A) Schematic of social exposure protocol of female mice with male mice over days. B) Scatter plot for comparison of mean response rates to the common first syllables in SN and SR, following exposure. C-D) Scatter plot to compare mean rate responses to common syllables (C) and disyllables (D) as in Fig. 2D and Fig. 3 respectively. E) Comparison of mean overall selectivity to sequences in SN to that in SR and comparisons across groups before exposure (AwF-BFEx, AF-BFEx and AM-BFEx) and after exposure (AF-AFEx). F) The mean selectivity to SN and SR in AF-BFEx in E) are compared with selectivity to SN and SR over days of exposure and within days of exposure. Violin plots for each day show the variability of the data sets. Thick green line shows percentage of units with higher selectivity to SN than to SR (axis to the right).

Thus we hypothesize that the entire sequences in SN are treated as objects that are behaviorally relevant in the male female exposure. Further, sequences in SNs appeared as the most informative set of sequences in the exposure event, over multiple days. Thus we compared rate based sequence selectivity of individual neurons (see Methods) and compare mean selectivity to SR and SN in the population of neurons in each group (Fig. 4E). Both AwF-BFEx and AF-BFEx showed higher selectivity to SN than SR (*paired-ttest*, p<0.001) with no difference between the two groups of females (*ANOVA,* p>0.05, SN, p>0.05, SR), reiterating lack of difference in such selectivity in the awake and anaesthetized conditions. AM-BFEx also showed increased selectivity to SN compared to SR, and surprisingly, had higher selectivity for SN than naïve females (*ANOVA*, p<0.001, AwF-BFEx, p<0.001, AF-AFEx). The AF-AFEx group of mice showed the highest difference between selectivity to SN and to SR. Interestingly, females after exposure had lower selectivity to SR compared to all other groups (*ANOVA* p<0.001, AwF-BFEx, p<0.001, AF-BFEx, p<0.001, AM-BFEx). Also, the selectivity to SN of AF-AFEx was similar to that of AM-BFEx (*ANOVA*, p>0.05) but significantly higher than naïve females (*ANOVA*, p<0.001, AF-BFEx, p<0.001, AwF-BFEx).

To further test that the increased difference in selectivity in experienced female mice was due to the exposure to male mice we considered the effect of multiple days of exposure (D1, D2, D3 and D>3, Fig. 4A). First we observe that on every time point considered, selectivity to SN was significantly higher than selectivity to SR during the exposure period (Fig. 4F,*paired t-test*, p<0.0001, D1, p<0.0001, D2,p<0.0001,D>3,p<0.0001) The effect of exposure over days shows that the selectivity to SN in AF-AFEx was significantly higher than that in AF-BFEx on all the days (*ANOVA*, p<0.0001, D1, p<0.0001, D2, p<0.001, D3, p<0.0001,D>3). Moreover we found that selectivity to SN increased significantly over the first two days (*ANOVA,* D1>BFEx, p<0.0001, D2>D1, p<0.0001) following D2, there was no significant change in selectivity to SN and was saturated. On the other hand selectivity to SR was unchanged on D1 (*ANOVA* p>0.05) and decreased on D2 (*ANOVA*, p<0.0001) and then remained unchanged over further days (*ANOVA*,p>0.05, D2/D3, p>0.05, D3/D>3) while being significantly less than that of naïve females and that of AF-AFExD1 (*ANOVA*, p>0.05). These observations are clearly consistent with the idea that exposure or experience dependent plasticity occurs to increase selectivity to the entire sequence but not its component syllables and disyllables. Since responses of SR and SN are from the same neurons collected in pseudo random order, suggests that neurons with higher selectivity to SN also lose selectivity to SR over days to maximize differences of the relevant sequences from any others. Thus although the AFExD2 group of mice were never exposed to the SR we find a decrease in selectivity to SR. The difference in selectivity to SN and SR saturates after D2 and it is explained by the fraction of units in L2/3 of A1 that have higher selectivity to SN than to SR (Fig. 4F, green thick line, *chi-square test*, p>0.01). Fraction of units with higher selectivity for SN than SR significantly increases on D2 from D1 (*chi-square test*, p<0.05) coincident with the decrease in mean selectivity of SR.

### Somatostatin (SOM) positive interneurons and Thy-1 excitatory neurons behave differentially during observed experience dependent plasticity in natural sequence selectivity

To elucidate the mechanisms underlying the observed plasticity, we hypothesized the involvement of inhibitory interneurons as observed in multiple studies^54^. Our single unit recordings are mostly regular spiking and hence extracting putative fast spiking inhibitory neurons is not possible. Our final conclusions above are based on responses to entire sequences (0.389s – 1.232s) and not fine time scale activity information. Thus Ca^2+^ dependent fluorescence imaging was performed in naïve and experienced female mice in the awake state through a cranial window (see Methods). We used 3 types of mice to image activity of Thy-1 positive excitatory neurons (EXNs, JAX- 24339, n= 3 mice) and genetically identified individual inhibitory neurons types (INNs, SOM JAX-13044 crossed with JAX 24105, n= 3 mice and parvalbumin positive, PV, JAX-8069 crossed with JAX-24105, n= 2 mice).

We first performed widefield Ca^2+^ imaging^55,56^ (see Methods) to identify location of A1 with UF ^57,58^ and other auditory areas^59^ (Fig. 5A). Following identification of A1 (see Methods), fine scale 2- photon imaging (see Methods) with single neuron resolution was performed by restricting individual ROIs (120-150 um x 300-350 um) within A1. Single ROIs (Fig. 5B, 2-3 per day) were imaged in each group of mice, female before exposure (AwFIm-BFEx, Thy-GCamp, Fig. 5B *left*, SOM-GCamp, Fig. 5B *middle*, PV-GCamp, Fig. 5B *right*) and after exposure (AwFIm-AFEx. Chronic recordings in this case allowed collecting data from same animals over different days of exposure (5 days), however, same neurons could not always be tracked reliably over days. As expected from the literature we obtained data from 100+/-20 neurons in each ROI of Thy-GCamp mice and 6+/-4 and 8+/-4 neurons in each ROI from SOM-GCamp and PV-GCamp mice respectively. Within an ROI, many neurons produced significant responses (mean df/f compared to mean baseline df/f, *ttest*, p<0.05, see Methods) to one or more stimuli in all the cases (Fig. 5C).

**Figure 5.**
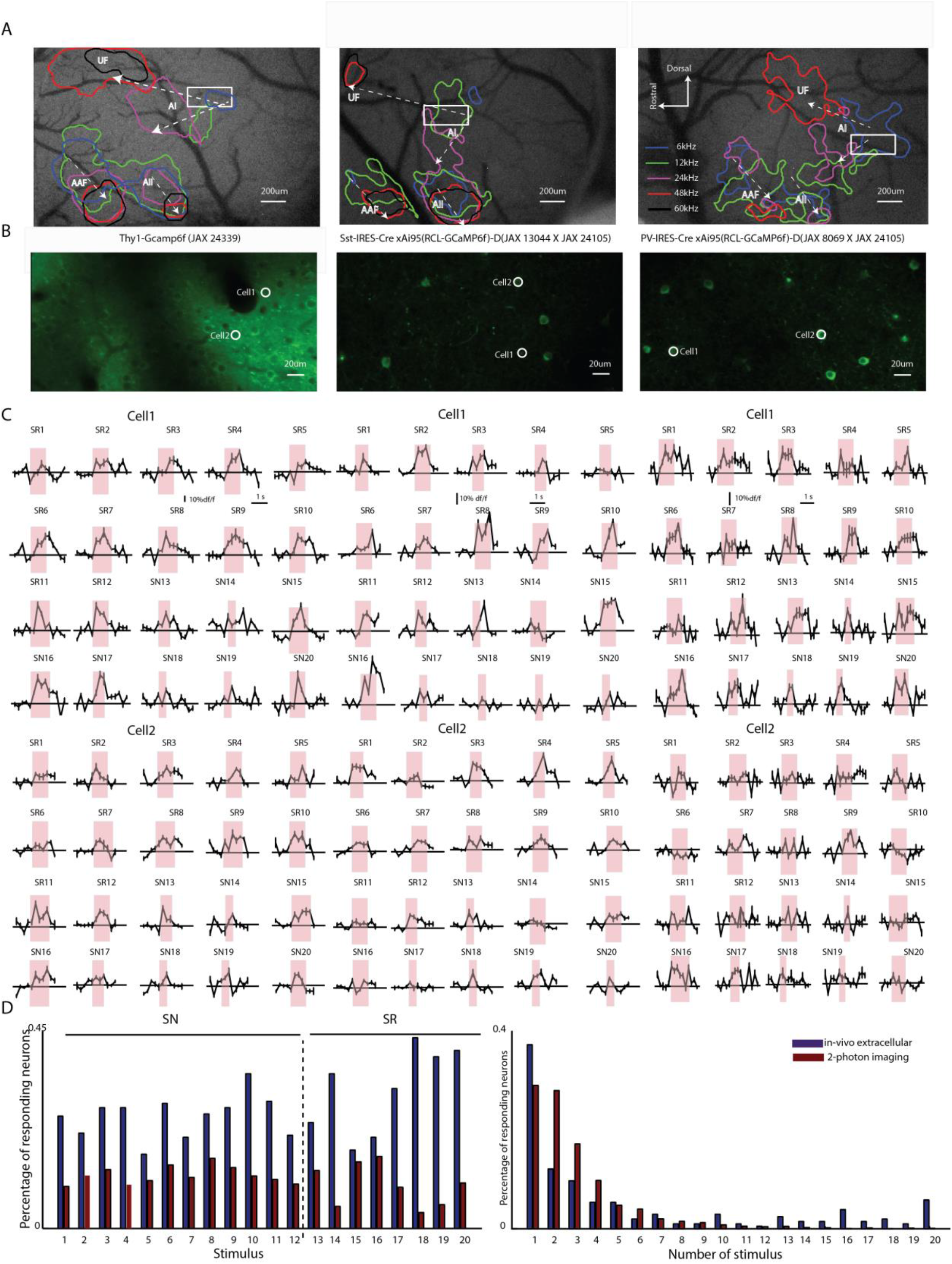
Sparse responses to sequences obtained with two-photon Ca^2+^imaging of Thy1, SOM and PV neurons. Representative examples of tonotopy in A1 and other auditory areas in mouse A1 obtained in all 3 groups of mice (Thy- 1-GCamp, SOM-GCamp and PV-GCamp in 3 respective columns) are shown. White box marks the area shown in B) below. Sample 2-photon image of an ROI in A1 in each of the 3 groups of mice. C) Average df/f plots obtained with 2-photon imaging, in response to each of the 20 stimuli for 2 cells in each ROI (of B marked as Cell 1 and Cell 2 in each column). D) *Left*: Bar graphs show percentage of single units (blue) with significant rate responses to each stimulus (1-20) in AwF-BFEx and that of single Thy-1 positive EXNs (brown) with significant responses in Ca^2+^ in AwF-BFEx. *Right*: Bar graphs show (same color representation as in *left*) percentage of neurons responding to either 1, 2, … or all 20 of the stimuli.

Distinct differences in response behaviour to sequences existed in population of neurons between single units (primarily, EXNs) and Thy-1 neurons’ Ca^2+^ responses. In general responses are sparser with 2-P imaging and underlie some of the discrepancies observed between imaging and extracellular recordings^58,60,61^. Fraction of neurons responding to each of the 20 sequences (SN and SR) in the awake state show lower fractions with imaging compared to that with single unit recordings in the awake state (Fig. 5D). Similarly fraction of neurons (above two groups) responding to the number of the 20 stimuli showed that the single unit population responded less selectively than the population of neurons observed with imaging (means 2.9 and 5.6 stimuli respectively with Ca^2+^ imaging and single units, Fig. 5D, *chi-square test,* p<0.001). Thus direct comparisons with single unit data may not be possible when using Ca^2+^ based responses.

We considered the selectivity of the three different types of neurons to SN and SR and effect of exposure in two ways. First we considered the relative selectivity to SN and SR, by grouping the neuronal populations as to how many of the SR and SN neurons of each type respond simultaneously. We first observe that before exposure, A1 single Thy-1 EXNs and SOM INNs, respond to subsets of both SN and SR similarly, with neurons responding to few or no SR also responding to few or none of SN. Neurons responding to more SR are more likely to respond to more of the SN (Fig. 6A). However, with exposure, EXNs reduce the number of SN to which they respond (see histograms below, Fig, 6A). The same is true of SOM INNs but to a lesser degree. PV INNs on the other hand (Fig. 6A, *right*) show similar degree of responding simultaneously to SN and SR before and after exposure. Thus, it is likely that PV neurons are less involved in the observed exposure based change in selectivity to SN.

**Figure 6.**
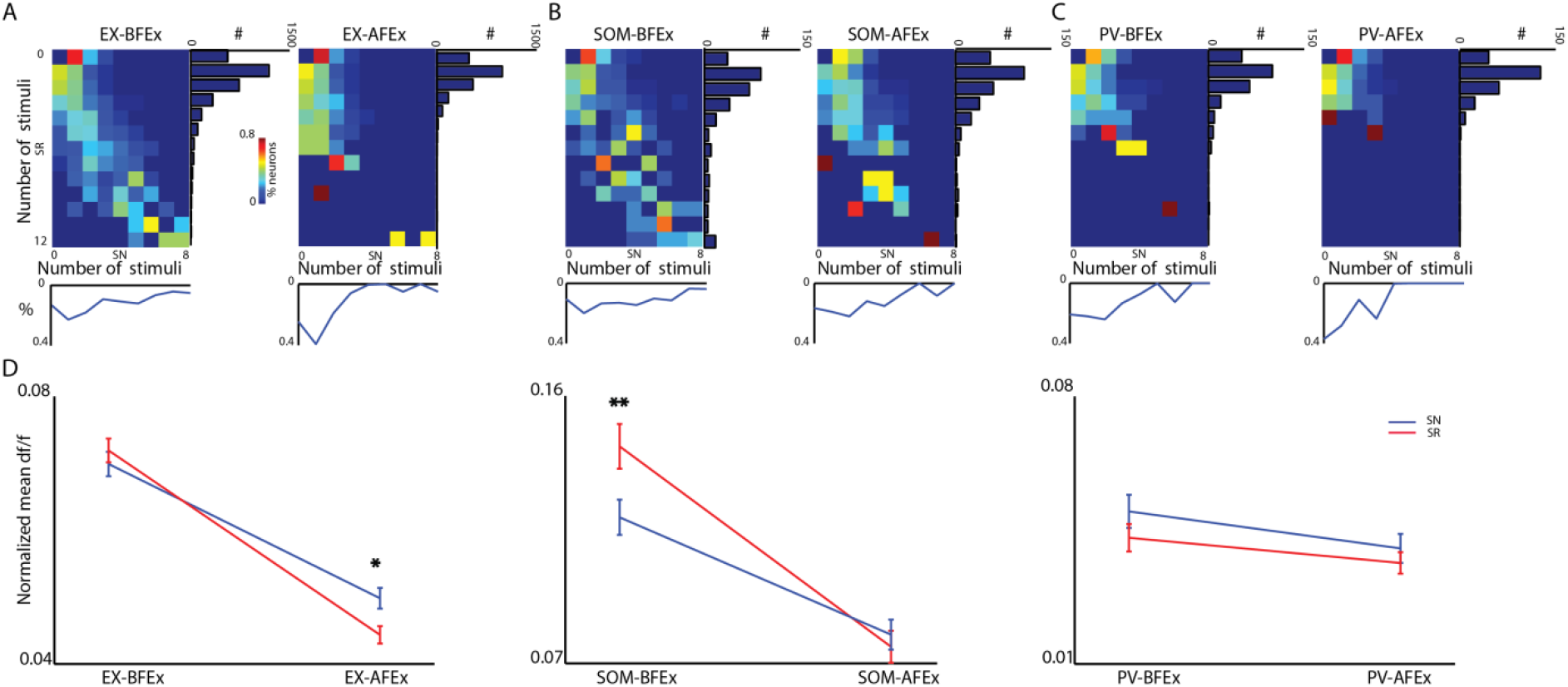
Differential effects of social experience driven plasticity in EXNs and SOM INNs. A-C) Each panel represents population data with 2-photon imaging in Thy1-GCamp (A), SOM-GCamp (B) and PV-GCamp (C) mice, with two matrix plots for before (*left*) and after (*right*) exposure. The rows in each matrix represent the percentage of neurons in the group and condition that respond to none (0), one, two … to all (8) of the SN (x-axis), of the neurons that respond to none (0), one, two … to all (12) of the SR (y-axis). Marginal distributions to the right show the number of neurons responding to the different number of SR. Distribution at the bottom of each matrix plot shows the average of the rows. D). Comparison of average selectivity to SN and SR in each condition BFEx and AFEx (before and after exposure) and between the conditions, for each neuronal type (Thy-1+, *left*, SOM+, *middle* and PV+, *right*).

Next we compared mean selectivity to SN and SR, based on responses quantified with df/f (see Methods) before and after exposure in Thy-1-GCamp female mice. We found that EXNs had increased selectivity to SN compared to SR (*ANOVA*, p<0.001) following exposure. Thus the observed experience dependent plasticity of entire SN sequences observed with single units was also observed with Ca^2+^ responses, relative to responses to SR (Fig. 6B *left*). However there was an overall decrease in response selectivity to both SR and SN, not observed in single units. The above departure from single unit population behaviour seen with Ca^2+^ imaging is due to the inherent differences in the two as stated previously (Fig. 5D). Thus with the Ca^2+^ imaging data we consider changes with respect to SR in each condition. Unlike EXNs, SOM INNs had significantly less selectivity to SR before exposure, which is abolished following exposure (Fig. 6B *middle*, *ANOVA* p<0.001). As with Thy-1 EXNs, exposure caused SOM INNs to also have decreased selectivity to both SN and SR. On the contrary, PV INNs neither showed a difference in selectivity to SN compared to SR both before and after exposure (*paired t-test*, p>0.05) nor did their overall selectivity vary with exposure (Fig. 6B *right*, *ANOVA*, p>0.05 SN, p>0.05 SR). Thus among the two inhibitory neurons tested, we hypothesize SOM INNs to be involved in the observed experience dependent plasticity. Also, overall SOM INNs had the highest selectivity both before and after exposure and thus are likely capable of mediating changes to specific stimuli.

### Optogenetic silencing of SOM INNs with sequence presentation induces plasticity in selectivity to sequences but not sequence components

To test the hypothesis of involvement of SOM in the above experience dependent plasticity of entire sequences we performed experiments in naïve anaesthetized female mice. Since AwFBFEx and AF-BFEx mice did not show any difference in sequence selectivity between them both for SN and SR use of anaesthetized mice is justified. Mice, (P56-P90) expressing ArchT-EGFP specifically in SOM neurons, obtained cross breeding of JAX-21188(Ai40D) and JAX-13044 (SST-IRES-Cre) were used (Fig. 7A). First we decided the power level of the 589 nm laser for ArchT-EGFP activation to silence SOM INNs. We used the highest power level at which the spontaneous activity post-light off recovered to initial spontaneous activity pre-light on, while producing an average increase in spontaneous with light on in the pre-stimulus period (Fig. 7A, *n*=6 mice,(3 female, 3 male) *n* = 51 units, noise at 5 intensities(55-95db SPL), see Methods^62^. There was no significant difference in spontaneous activity between pre-light on and post-light off periods (Fig. 7B *right*, *paired ttest*, p>0.05), with increased light-on-spontaneous activity (Fig. 7A, green box, Fig. 7B *left* and *middle*).

**Figure 7.**
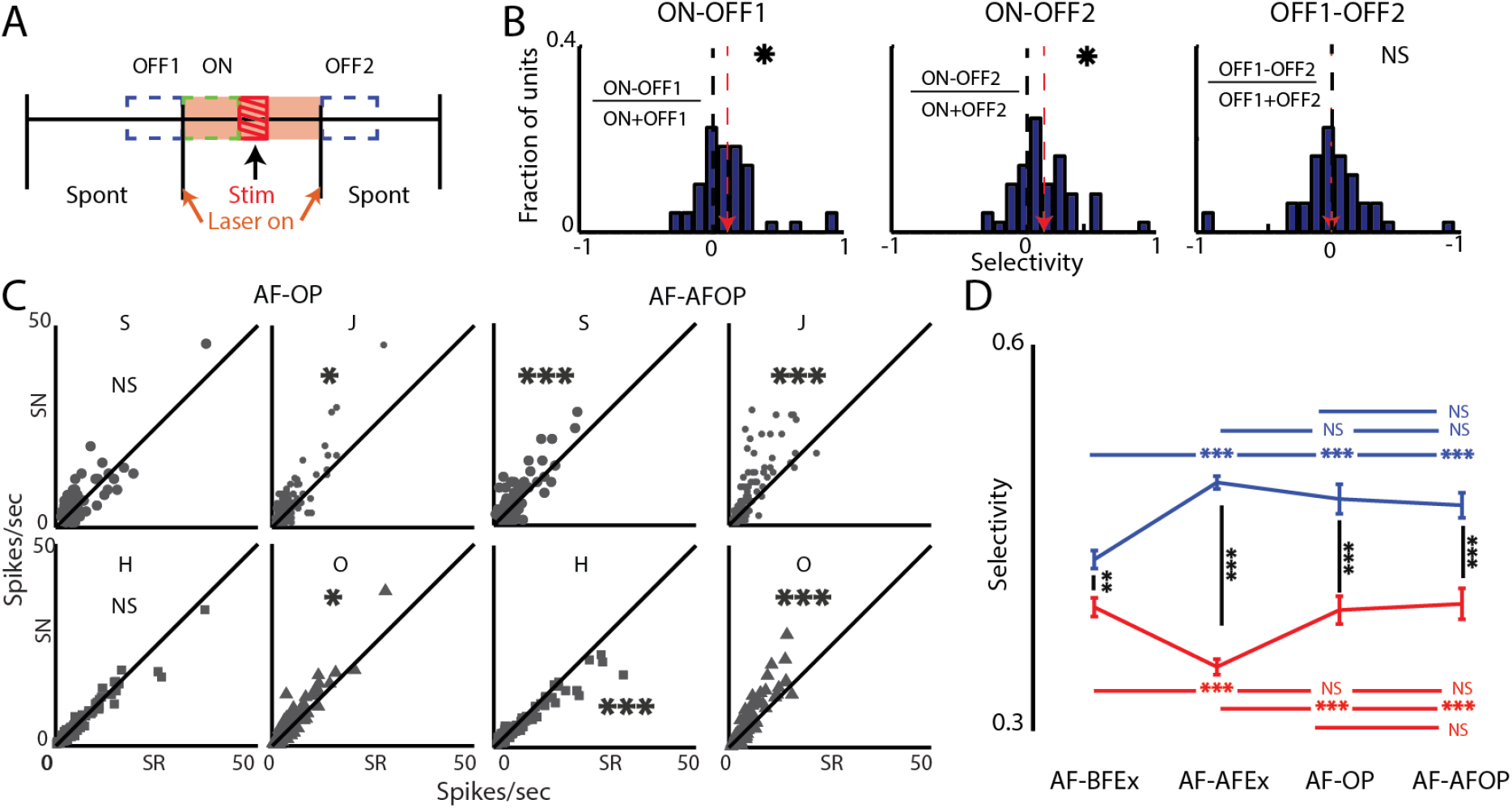
Reversible silencing of SOM paired with sequence presentation mimic plasticity in sequence selectivity without altering syllable selectivity. A) Schematic of sequence of events for optogenetic silencing of SOM and determining laser power. For all such experiments the laser was turned on 100 ms pre stimulus onset and turned off 100 ms stimulus offset (orange shading). Spontaneous or non-auditory driven activity in 3 periods were used, OFF1, ON and OFF2, each 100 ms long and were as depicted. For sequences stimulus onset and offset were onset of first syllable and offset of the last syllable of the sequence. B) Histograms of modulation index of all cases show significant modulation of spontaneous spiking by light (*left* and *middle*, mean: red arrow), and the histogram to the *right* shows comparisons of spontaneous activity pre and post light on. C) Scatter plot to compare single syllable mean responses in SN and SR during SOM silencing paired with sequences (AF-OP) and after the period of pairing (AF-AFOP), as in Fig. 2D and Fig. 4C). D) Similar plot as in Fig. 4E, with AF-BFEx, AF-AFEx (from Fig. 4E) and additionally AF-OP and AF-AFOP.

We performed optogenetic silencing of SOM INNs at the above power over a period encompassing presentation of the 20 sequences (both SN and SR, 100 ms before to 100 ms after the stimulus as in Fig. 7A, see Methods) for total 100 presentations (5 for each SN and SR) to test our hypothesis of involvement of SOM neurons. We found that the relationship of responses of single syllables in SN and SR during SOM silencing (AF-OP, Fig. 7C, *left*) were similar to AF-BFEx except the reduced responses of H in SN compared to SR was now the same (Fig. 2C, *middle*). We also probed for longer term plastic changes induced by the pairing of SOM INN turning off (effectively disinhibiting EXNs to which they primarily project)^36^ and simultaneous sound stimulus (SN and SR) presentations. We found that the comparative responses to each of the single syllables in SN and SR following the above pairing (AF-AFOP, see Methods) were largely the same as AF-BFEx, AwF-BFEx and AF-AFEx (Fig. 2C and Fig. 4C, Fig. 7C, *right*). No distinct switch in preference between SN and SR was observed. Similar observations were made with transitions common to SN and SR in female mice before and after exposure (Fig. 3 BFEx, Fig. 4D AFEx) and during and after optogenetic silencing of SOM INNs (Fig. S5), with no distinct switching between SN and SR. We then tested the idea of plasticity to selectivity in sequences. For both groups AF-OP and AF-AFOP, there was a significant increase in selectivity to SN compared to the AF-BFEx group and was same as that of the AF-AFEx group of mice (Fig. 7D, *ANOVA*,p<0.0001, p<0.0001, p<0.0001). The mean selectivity to SR remained the same as the AF-BFEx group. In the subgroup of units from AF-BFEx mice in which SOM INN silencing based pairing was performed, the mean selectivity of SN was the same as AF-BFEx however there was a small reduction in selectivity to SR in the same population (median 4 % lower) indicating an effect of rise in selectivity to SR during and after pairing. Thus SOM inhibition during sequence presentation is capable of inducing rapid plastic changes in coding of entire sequences without changing coding of the component syllables and disyllables. We thus hypothesize that during the activity dependent plasticity occurring in naïve females, VIP mediated inhibition of SOM could underlie the plasticity in sequence selectivity.

## Discussion

Our work is the first study that investigates selectivity of A1 single neurons to sound sequences as a whole and not only by its component tokens and dyads^29,35,63,64,65^. We also find experience dependent plasticity in selectivity to mouse vocalization sequences as a whole, unlike what is observed with maternal plasticity in encoding of single syllables of mouse pup calls^29^. Multiple studies have shown context specific vocalizations in adult male mice^11,13,39^. Variation in USVs in different contexts have primarily been characterized in terms of spectrotemporal content and basic call features like frequency of types of calls, duration and length of bouts among other such features 66,12. However, A1 coding of entire sequences are lacking, although, structure in call sequences in adult male mice ^39,43,46^ and pups^42,47^ have been known to exist.

Multiple methods have been used to evoke adult male USVs relevant for social interaction with females^11^. We develop a sequence of steps in male female interaction over days (Fig. 1A) from which we extract male USV sequences that are highly informative using information theoretic methods. Although in the above interactions females may also vocalize their contribution is minimal^67,68^ and primarily with low frequency harmonics^69,70^ not considered in our data. The main factor that warrant further investigation of whole sequence coding are our observation of significantly higher preference of female mice for the above extracted SN over the designed SR (Fig. 1). The SR was designed to have the same frequency of occurrence of each syllable type as found in our male female interaction protocol (Fig. 1C). Both SN and SR were constructed with the exact same syllable waveforms and repetition rate, removing any spectrotemporal cues other than syllable order to differentiate SN and SR. Control experiments with number of syllables in SR to match that of SN also showed the same results as a fixed number of syllables in SR, in terms of preference of females for SN (Fig. 1F).

The sequences extracted and used in our study are based on predictability from syllable to syllable. A set of such syllables combined into structured sequences as above, is capable of providing more information than the same syllables produced in isolation^20,^. In many species like the songbird^2^ acoustic communication takes the form of ordered sequences especially with more flexible vocal production. Sequences contribute to information-rich communication, as is the case for human speech as studied through prediction of letters in English text^20^. Recent work with machine learning have been able to connect single mouse USV syllables with particular behaviour^71^ and also extracting information about identity, sex and context^72^. However, similar approaches have not been taken for mouse USV sequences of syllables. While we do not claim that each sequence may convey different meaning or information, but the results do provide the fundamental bases required for sequence based communication. Future studies need to combine more refined behaviour to link sequences with behaviour and further understand encoding and plasticity of their representation at single neuron and network level.

Long term as well as rapid alteration of representation of sounds in mouse A1, with experience, like maternity ^28,29^, learning based on reward or fear^32,30^, developmental environment ^55,73,74^ and artificial involvement of subcortical and higher order structures^75,76^ have been shown for single tokens of natural as well as artificial sounds. However, our study shows social experience dependent change in selectivity to behaviourally relevant sequences as a whole and not of its parts (Figs. 2–6). Our results show that A1 responses of single tokens and dyads depend on context, but their relative selectivity do not change with the above experience (Figs. 2–6) while selectivity of single neurons for the whole sequence is altered based on the degree of the experience (Fig. 4F). Integration of acoustic components into a single percept across frequency and at least partially overlapping in time is well known ^30,34^. However feature integration to obtain a holistic representation of sound tokens over non-overlapping in time is surprising and not well understood. Such integration likely involves very long time scales of adaptation known in A1^77,78^, long time constant recursive connections and inhibitory inputs^79^. Previous work^52^ has looked into very long time scale adaptation ^78^ of entire sound sequences and change in their representation over time from repeated presentations and show recurrence in EXNs and SOM to play a role. Here we find that optogenetic silencing of SOM INNs paired with sequence presentations alter sequence selectivity as with social experience with largely no change in relative selectivity of single neurons to syllables.

We hypothesize that the above silencing of SOM INNs in naïve female mice during the social experience occurs through VIP neuron activation triggered by the interaction with the male. SOM INNs are known to disinhibit EXNs upon activation of VIP neurons^36^. SOM neurons respond less selectively to SN compared to SR before exposure and EXNs behave in the opposite manner. Thus higher responses to EXNs to SN compared to SR coupled with disinhibition by SOM can drive the observed plasticity. Higher selectivity to SR before exposure of SOM INNs would reduce the disinhibition when SR is presented and thus produce less plastic effects. However, activation of VIP during the said social experience is unknown and needs to be explored. The likely candidates are inputs from BLA(Basolateral Amygdala), hypothalamus, Ventral tegmental Area (VTA) or prefrontal cortex because of their involvement in mating behaviour^80, 81, 82^.

Our results take us a step further in the direction of establishing mouse models of vocal communication and to study disorders like ASDs and other neurobiological disorders with social communication deficits like intellectual disability and aphasia. Our study also invites reinvestigation of many aspects of mouse communication and even vocal learning ^9,10^ and established ideas of innateness of mouse USVs, by considering sequences of syllables.

## Methods

### Animals

All mice used in the study were 8–12 weeks old at the time of experiments. All procedures pertaining to animals used in the study were approved by the Institutional Animal Ethics Committee (IAEC) of the Indian Institute of Technology Kharagpur. Mice, *mus musculus* (age and sex identified in individual cases), are reared under a 12/12 h light/dark cycle at a temperature of 22–25°C with *ad libitum* access to food and water. All vocalization and extracellular recordings are done on C57BL6/J strain mice. For two-photon imaging, the PV-ires-Cre [JAX 08069] and SOM-ires-Cre [JAX 13044] driver lines are crossed Ai95(RCL-GCaMP6f)-D[JAX 24105] to label inhibitory neuronal types with green fluorescence expression. Recordings from excitatory neurons are done by using C57BL/6J-Tg(Thy1-GCaMP6f) [JAX 24339].

### USVs Recording

To record adult male mouse USVs, initially a male mouse was kept in a wooden recording cage (12×18×15 cm) placed in a sound isolation booth (Industrial Acoustics, New York, NY) alone for 5-10 minutes. No vocalizations were observed in the above condition. A female was introduced in the same setup with a separator in between (5-10 minutes), followed by removal of the mesh in between them. USVs were emitted by the male in both the latter cases. After 7-10 days, when the male was placed alone in the recording set up, it vocalized in the absence of the female. Final recordings were made starting from the above stage and analysed further. USV recordings were collected using the protocol depicted in Figure 1(A). Context1: The male mouse alone (M); Context2: The male mouse with a female present in view, separated by a mesh (MSF); Context3: As context2 without the separator. The above context specific USV recordings were made for at least 5 days. Acoustic signals were recorded with a free-field microphone and amplifier (1/4” microphone, Model 4939, Bruel and Kjaer, Naerum, Denmark) with flat frequency response up to 100 kHz, and slightly diminishing sensitivity at higher frequencies. The acoustic signals were digitized at 250 kHz with 16-bit resolution collected with National Instruments DAQ. Recorded signal time waveforms and spectrograms were displayed in real-time on a computer with open access bioacoustics software, Ishmael.

### Vocalization Recording Analyses

All our USV analyses were performed in MATLAB (Mathworks) and have been presented in detail earlier^42^. In brief, each wav file was divided into 5 second epochs. Background noise was eliminated bandpass filtering with a Butterworth of order 8, removing frequencies outside the 20kHz and 120kHz range. Syllable segmentation was performed by first calculating the Short Term Fourier Transform (STFT) of each epoch with 1024 length Hamming window, overlap of 75%. Syllables were identified by calculating the power concentrated in each frame normalized by the average power in all the frames and median filtered over 30 ms windows. Peaks in power over time were detected by using peak detection. Syllables were classified into 5 categories^42^, Noisy (N-type) syllables which has a broad energy content, syllables with harmonic content (H-type), based on the presence and absence of discontinuities, syllables are again classified into three classes, namely S-type (continuous contour of spectral energy), Jump, J-type (a single discontinuity) and, Other, O-type (more than one discontinuity)(Fig. 1B). All subtypes of syllables are also present, as observed in other studies^12,13,83^. However, we used only pitch jump as the primary classification criteria based on Holy and Guo^43^ to restrict the number of broad classes enabling our analyses requiring large sample sizes. Moreover, sudden discontinuities in pitch are also inherently tied to the vocalization production machinery.

### Identification of high probability sequences in different contexts

The significance of occurrence of each syllable at different positions given the previous syllables was computed for each context. For the syllable in the first position in a bout, the probability of occurrence of each syllable type in the beginning of the bout was obtained and compared to the equally likely probability of occurrence of each syllable type. Syllables with higher (90% confidence) probability of occurrence than overall were considered as significant. The process was continued for each of the subsequent positions keeping the previous syllable types fixed until there were no significant syllables further observed. In the above manner we find sequences that occur above chance and render structure in different contexts.

### Surprise Analysis

Surprise was computed by calculating the dissimilarity between the posterior and prior distributions of occurrence of syllables in sequences (length 3 - 7) using KLD^44^ as the sequences used as stimuli had minimum 3 and maximum 7 syllables. Surprise was defined by the average of the log-odd ratio^48^

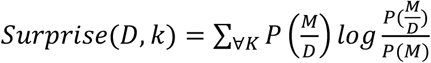

where *P (M*), the prior probability *(D)*, the posterior probability and *k*, the space model of the sequences.

### Sequence Construction and Auditory Stimulus Delivery

The stimulus consisted of 12 random sequences drawn randomly based on the probability distribution of syllables (Fig. 1C) obtained in 3 contexts (4 sequences from each context). The remaining eight sequences were the high probability, high surprise natural sequences emitted in different contexts. Between each syllable, a silence of 90 ms was considered based on the mean of the ISS distribution (Fig. S1C). All stimuli are generated using custom-written software in MATLAB (Math-works) and D/A board (National Instruments), attenuated using a TDT PA5 (Tucker Davis Technologies, TDT), generated using TDT EC1 calibrated speakers (driven with TDT ED1 drivers), and delivered through a speaker kept 10 cm away from the contralateral ear. The acoustic calibrations, performed with microphone 4939 (Brüel&Kjær), of the ES1 speakers (TDT) in the sound chamber, showed a typical flat (95 dB) calibration curve from 4 to 60 kHz; 0-dB attenuation on PA5. Each of the syllables in the sequence had an onset and offset of 5ms ramp and had root mean square (rms) matched. All awake and anesthetized recordings were done at 65-75 dB SPL and 75-85 dB SPL respectively. Responses with usually 5 (rarely 4 or 6) repetitions were presented in randomized order and used for further analysis. Each stimulus onset was preceded by a baseline of 500ms and had an inter trial interval of 5 s.

### Two choice free access/preference task: Natural vs Random

All behavioural tests were performed inside a soundproof anechoic chamber under dim red light. The test cage was an acrylic rectangular box (30-cm long × 20.5-cm wide × 20-cm tall). Two tubular ports (9 cm long) terminated with a mesh on one end, each 5cm in diameter, were attached to the test cage opposite to each other, one to the right corner (RS) and the other to the left corner (LS) (schematic of the set up shown in Fig. 1F) with a speaker beyond the mesh end of the ports. Before starting a session, the entire test cage was wiped with 70% ethanol. There were five sessions each of 5 min, first 4 of them are shown in Fig. 1F (S1 to S4). In S1 the test female mouse was allowed to explore the test cage; in S2 SN (say from RS) and SR (from LS) were played alternatively with an inter stimuli gap of 5s; in S3, no stimulus; S4 was same as S2 with corners swapped for SN (from LS) and SR (from RS); in the final session again no stimulus was played. A webcam (Logitech C925e) was fitted 35 cm above the center of the test cage from the base and the entire duration of the 5 sessions were recorded at 15 frames/second, 1411 kbps, using Logitech software.

### Calculation of Joint Distributions and *MI* Based Dependence

Transition probabilities were computed by estimating the joint probability distribution. Krischevsky and Trofimov (KT) correction was applied to take care of combinations leading to ‘0’ values. Mutual Information or MI, between 2 random variables X and Y, quantifies total dependence between the two random variables^44^ and can be computed from the joint distribution *P(X, Y)* and its marginal distributions *P*(*X*) and *P*(*Y*) as

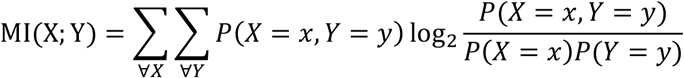

Since MI were sensitive to bias^45,84^, the results can be erroneous provided the limited data size. The above problem was circumvented through bootstrap removal of bias and comparing the *MI* estimates with scrambled syllable order in sequences of vocalizations to get only significant estimates of *MI*.

### Removal of Bias in Mutual Information Estimates and Significance Analysis

Bootstrap debiasing was used to remove bias in MI estimates^42,45^. A resampling technique where the bootstrap dataset has the same number of elements as the original one was employed to debias the estimates get an estimate of the bias perform debiasing and obtain confidence intervals. We compare the lower confidence interval with upper confidence interval of the estimate of ‘0’ *MI* obtained similarly as above but now by not keeping the transitions intact, that is by randomly scrambling the sequences which leads to the estimates of ‘0’ *MI* and its confidence interval from the same data set with same number of syllables and other statistics intact except the order of the syllables in a sequence. When the confidence intervals of *MI* of the data and the scrambled data did not overlap, the estimated *MI* was considered to be significant. Thus we minimize the possibility (<5%) of spurious *MI* due to limited data size and variability.

### Calculation of Kullback Leibler Divergence (*KLD*) between distributions

The proximity among the probability distributions was quantified using *KLD* ^44,45^, an information theoretic distance metric which makes no assumptions about the statistics of the data. To compute *KLD* between the distributions *P* and *Q* taking on values over the same set (in our case syllables produced by three different contexts taking values of different syllable types with probabilities *P*(*x*) and *Q*(*x*), *x* being a syllable type, or the syllable to syllable transitions produced by the two groups) was computed as follows:

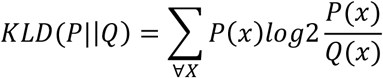

We performed debiasing of *KLD* using bootstrap resampling calculated significance in the same way with 95% confidence intervals as with MI (above).

### Tracking Mouse Movement

The videos of the 5 sessions monitoring the behaviour of mice in the syllable sequence preference test were analysed using custom code written in MATLAB. We used the MATLAB computer vision toolbox function ‘*kalmanFilterForTracking*’ for tracking the position of the mouse in the test chamber in every video frame during the 5 sessions. When the mouse was not visible in the chamber, when the mouse was inside a port, the position of the mouse was assigned to the last observed location, always near the entry point of the port. If in the initial frames the mouse was not visible, the mouse location corresponding to all such initial frames was assigned to the co-ordinates of the position the mouse was first detected. A bounded tracking was done, by selecting a region of interest covering the entire test floor and the entire area outside it was never assigned as a possible mouse location. Based on the manually selected co-ordinates of the four corners of test chamber the area was then divided either into 2 equal halves corresponding to RS and LS respectively or 3 equal parts corresponding to RS, LS and a central neutral region. Based on the tracked co-ordinates of the mouse the percentage of time spent towards each corner (RS or LS) was quantified for the different sessions and used for statistical comparisons.

### Surgical Procedure for in vivo recordings

The mouse is anesthetized in an induction chamber with isoflurane. It is placed on a heating pad (for maintaining body temperature), and isoflurane inhalation is maintained via a nose mask. An incision is made in the scalp along the midline. The region of interest is drilled using a micro drill, and an electrode is implanted. It is cemented using a super bond. The animal is habituated for 5-7 days. The recording is done in a single-walled sound-attenuating chamber. On experimental days, the animal is placed securely into a foam body mold. The headpost is attached to a custom-made stereotaxic apparatus. For imaging, initially the mouse is injected with 0.1 ml dexamethasone on the thigh. Once the animal is stabilized on anaesthesia, hair remover cream is applied to the area of interest. After leaving it for a few minutes, it is cleaned. The remaining procedure remains the same as described above. For 2 photon imaging the craniotomy is sealed with a coverslip.

### In vivo extracellular recordings

Extracellular recordings were performed using a tungsten microelectrodes array (MEA) of impedance 3–5 Mohm (MicroProbes); 4×4 custom-designed metal MEAs with an inter electrode spacing of 125 mm were used. The array was advanced slowly to a depth of 200-350 um from the surface into the ACx using a micromanipulator (MP-285, Sutter Instrument Company). The electrodes were allowed to settle for 15-20 min before the stimulus presentation was started. Signals were acquired after passing through a unity gain head stage (Plexon, HST16o25) followed by PBX3 (Plexon) preamp with gain of 1000, to obtain the wideband signal [used to extract local field potential (LFP), 0.7 Hz to 6 kHz] and spike signals (150 Hz to 8 kHz) in parallel and acquired through National Instruments Data Acquisition Card (NI-PCI-6259) at 20-kHz sampling rate, controlled through custom-written MATLAB (MathWorks) routines. Data collection lasted for < 2 weeks, with ∼1-2 hour long sessions every day for the electrode implanted animals in the awake state. Units collected on each day from the implanted recording electrodes were considered as separate units. All sound tokens presented in all kinds of stimuli had 5-ms rise and fall times.

### Analysis of in vivo extracellular recordings

Spike sorting was done offline in custom-written MATLAB scripts. Data were baseline corrected and notch filtered (Butterworth fourth order) to reject any remnant power supply 50-Hz oscillations.). Single-unit spike times were obtained from the acquired spike channel data using threshold crossing and spike sorting with custom-written software in MATLAB. Mean spike rate was calculated by taking the mean of the neuronal firing over the stimulus duration +20ms. A neuron was considered to be significantly responding if the spontaneous activity (200ms before stimulus onset) was significantly different from the neuronal activity (unpaired t-test<0.05).

All the units which significantly responded for any syllable have been considered for calculating the neuronal selectivity. Mean spike rate is calculated for each of the 20 stimuli corresponding to their duration. For each neuron, then selectivity is calculated by

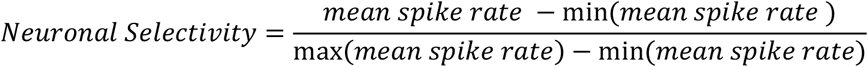

All the 8 natural stimuli are clubbed together by calculating their mean response across all neurons. Similarly, the mean neuronal selectivity is obtained for random sequences.

### Wide Field Calcium Imaging

After the animal preparation for chronic window implant, for wide field imaging, cortical images were taken using X-citeQ120 blue light (450-490 nm) excitation. A 4×0.13 NA objective (Olympus) was focused 200 microns below the cortical surface. The green emission fluorescence (500-550 nm) was collected onto a CCD camera at ~20 Hz.

For identifying the Auditory Cortex (ACx) regions, a significant change in fluorescence (df/f) was computed by subtracting and normalizing each frame by the mean response of the baseline frames. The pixels that have significant df/f retain their values (unpaired t-test). A Gaussian filter was used to smooth the df/f with a standard deviation of 2, normalized by the absolute maximum. A binary image is generated based on the pixels having values greater than 0.5. Using the inbuilt function of Matlab bwboundaries 8-connected neighborhood boundaries are drawn for different frequencies.

### Two- photon Calcium imaging

For two-photon imaging, Ultima IV, Prairie Technologies laser scanning microscope with a Spectra Physics Insight Ti-Sapphire mode-locked femto-second laser was used. Cells were imaged using a 20X, 0.8NA (Olympus) water immersion objective at depths of usually 200–350 μm from the cortical surface at an excitation wavelength of 860-920nm. Full frame images were acquired at a resolution of 512×512 pixels. The laser power was adjusted from 50mW to 80mW. Frames in the region of interests (120-150 μm x 300-350 μm with 1.16 μm pixel size) were imaged at ~4-10 Hz (~250ms frame period, 4 μs dwell time).

### Analysis of 2-photon imaging data

2-photon imaging analysis was performed using custom codes written in MATLAB (Mathworks). Imaging sequences were aligned by performing X-Y drift correction. Cells were selected manually by selecting the centre point of the cell on the motion-corrected mean images. ROI (5 µm radius) were drawn based on the cell centres. Raw fluorescence signals over time (f) of the selected ROIs across all frames were extracted. For each trial, relative fluorescence was computed by using df/f0 = (f–f0)/f0, where f0 corresponds to baseline fluorescence. Baseline fluorescence amplitude was estimated by calculating the mean of fluorescence values over all the frames preceding stimulus, except the first 3 frames, which was either 4 or 6 frames.

A neuron was considered responsive to a stimulus if the mean of three frames of the mean fluorescence trace (calculated from stimulus repetitions) before the stimulus onset is significantly different from moving three frame average after the stimulus onset (unpaired t-test, p<0.05). Baseline distribution was obtained from all the pre-stimulus 3 frames (all stimuli and all repetitions, usually 100). The total number of frames considered for each stimulus encompassed the stimulus duration and an additional 0.5s after the stimulus. We only considered significant positive going responses and thus FDR<2.5%. The response of a neuron to a sequence was calculated based on mean df/f of all frames from stimulus start to 0.5 ms post stimulus end. Selectivity to each of the 20 stimuli of a neuron was based on the above response and calculated as with single units.

### Calculation of selectivity based on spontaneous activity for optogenetics

For photo inhibition of SOM (JAX 21188XJAX 13044), we activated ArchT via light pulses of 589 nm^85^ through an optical fibre of 200 micron, 0.5 NA. The power at the tip is 30mW. Light pulses are presented 100 ms before auditory stimulus onset and last for stimulus duration+100 ms.

Reversibility due to photo inhibition of SOM is tested by comparing the mean spontaneous activity with (*W)* and without (*WO*) the laser being on. The formula used for calculating selectivity

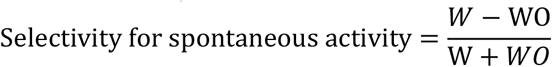

## Acknowledgements

SA thanks MHRD for Institute Fellowship, SB thanks India Alliance, IIT Kharagpur, MHRD and SRIC Cell, IIT KGP for start-up and Challenge Grant funds. *This work was supported by the DBT/Wellcome Trust India Alliance Fellowship/Grant IA/I/11/2500270 awarded to SB and India-Czech Bilateral Scientific and Technological Cooperation DST India.*

## Supplementary Material

**Figure S1.**
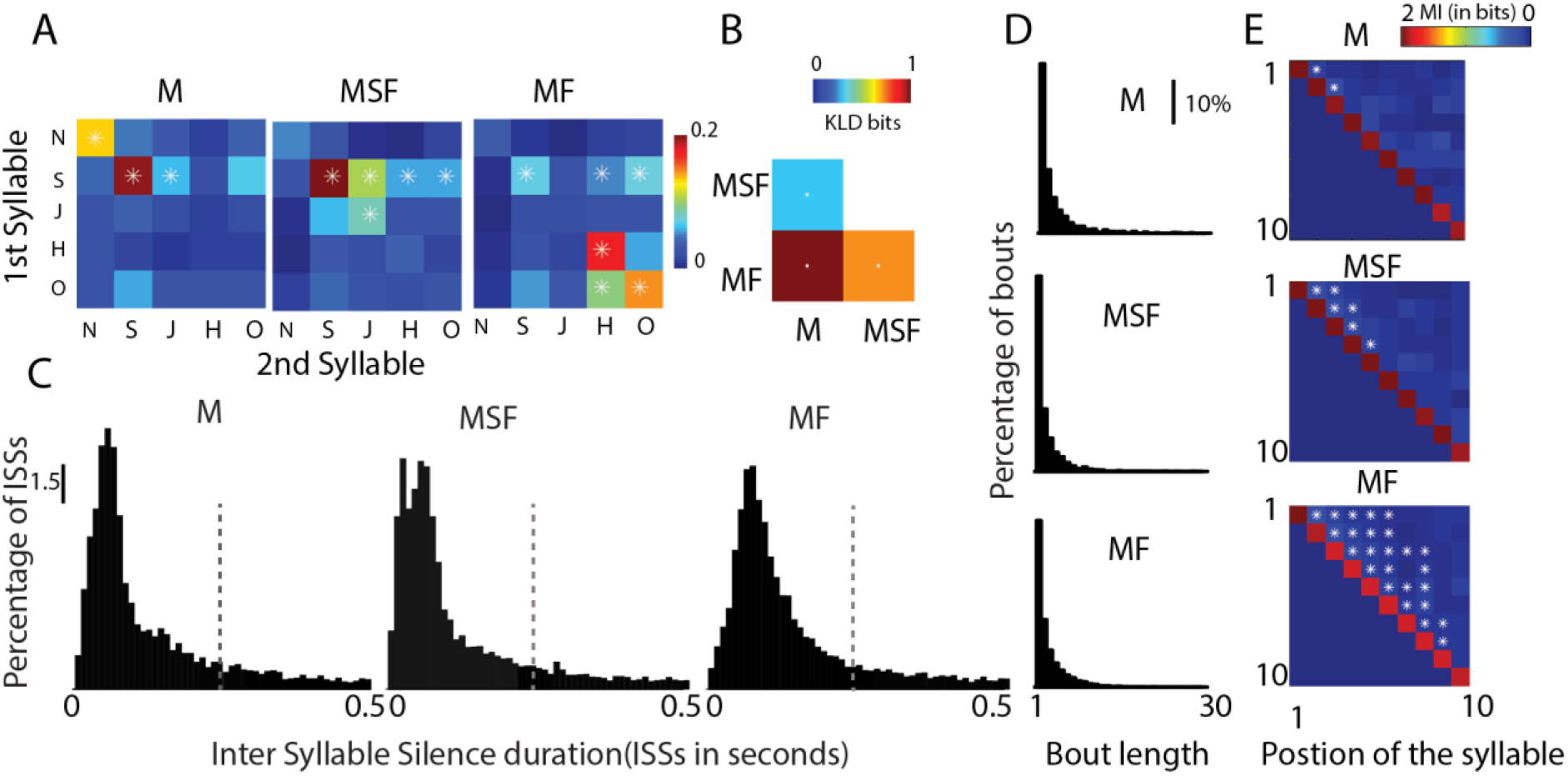
Context specific modulation in adult male mouse. A) Joint probability distributions of syllable to syllable transition considering starting 2 syllables in bouts is depicted in each of first 3 matrices in the row for the 3 contexts of the adult male (M, MF and MSF). B) The diagonal matrix quantifies the *KLD* between joint distributions shown in S1A. C) The three distributions depict the inter-syllable silences (ISSs) observed in each context of adult male. The vertical dashed line (at 250 ms) marks mean + 1*STD of the overall data. D) The distribution of percentage of bouts of a particular length present in each of the contexts is shown E) The 3 matrices represent the *MI* calculated as in Fig. 1D with each row showing the *MI* for the *n*^*th*^ syllable with the 1^st^ (row 1), 2^nd^ (row 2), 3^rd^ (row 3) and so on. The diagonal elements show the entropy of the syllable in the corresponding position from the bout start. Asterisks indicate significance at 95% confidence for (A), (B) and (D).

**Figure S2.**
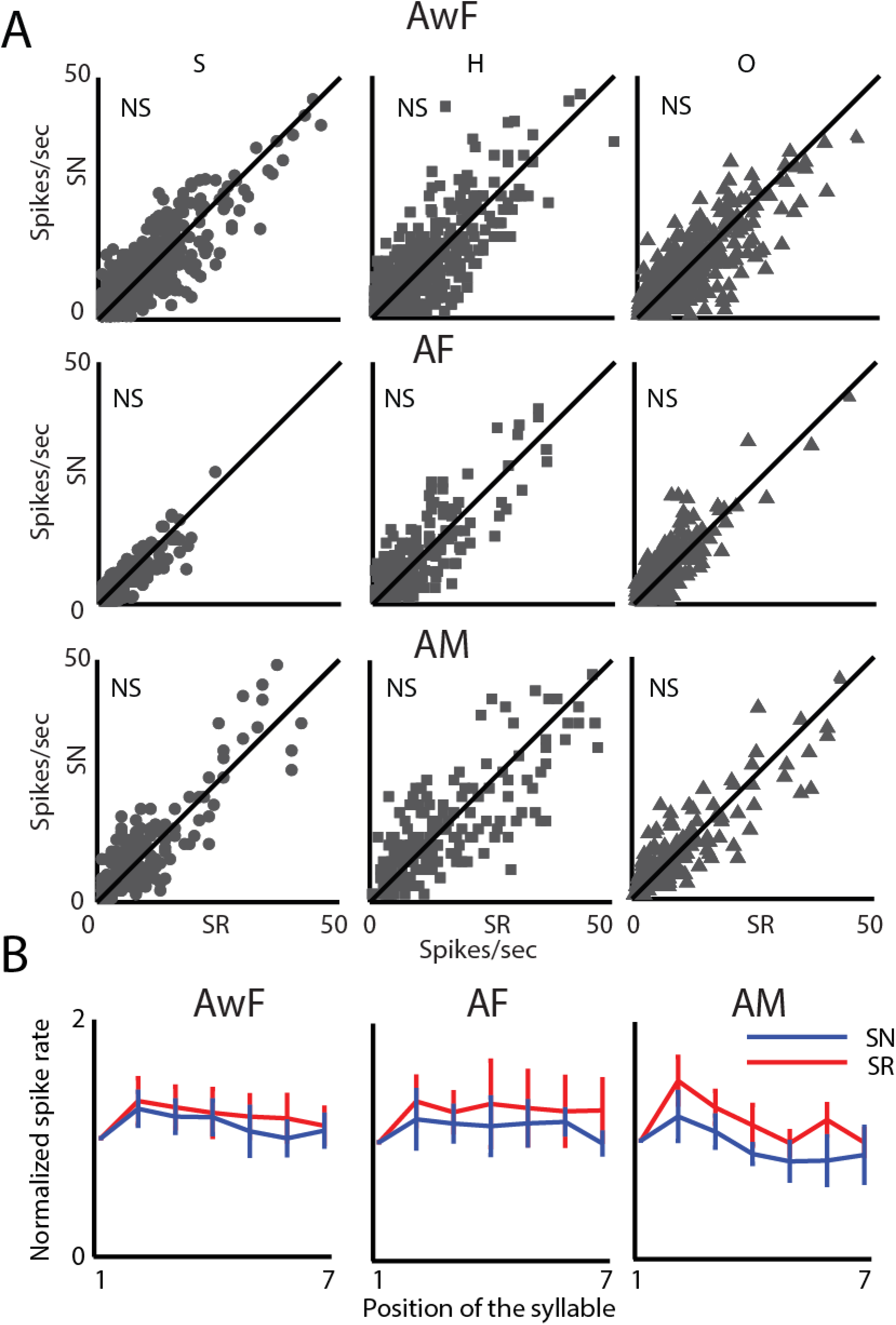
Similar mean response rate and time scales of adaptation for SN and SR. A) Scatter plots show comparison of mean response rates of all common first (identified by solid symbols, S: large circle, H: square, O: triangle) syllables types in SN and SR in 3 groups of mice, AwF, AF and AM. D) Profile of changes in the normalized response strength for each position of the sequences SN (blue) and SR (red) over time concerning the syllable in the starting position.

**Figure S3.**
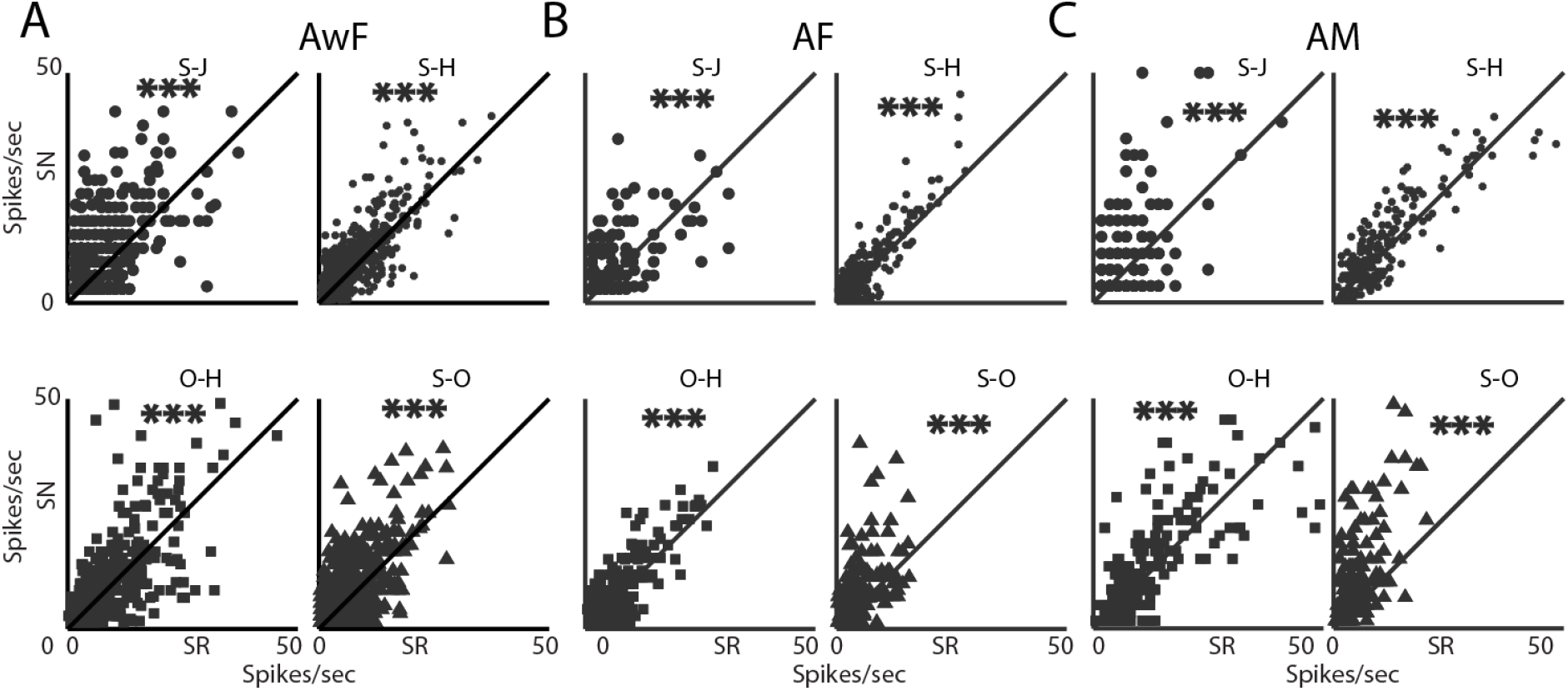
Context specificity in common disyllables with first transition in the starting position for SN. Scatter plot of comparison between mean rate responses to first transition in SN (S-J, S-H, S-O and O-H) and the same transition present at any position in SR based on response to the second component, excluding the first transition for the three groups AwF (A), AF (B) and AM (C).

**Figure S4.**
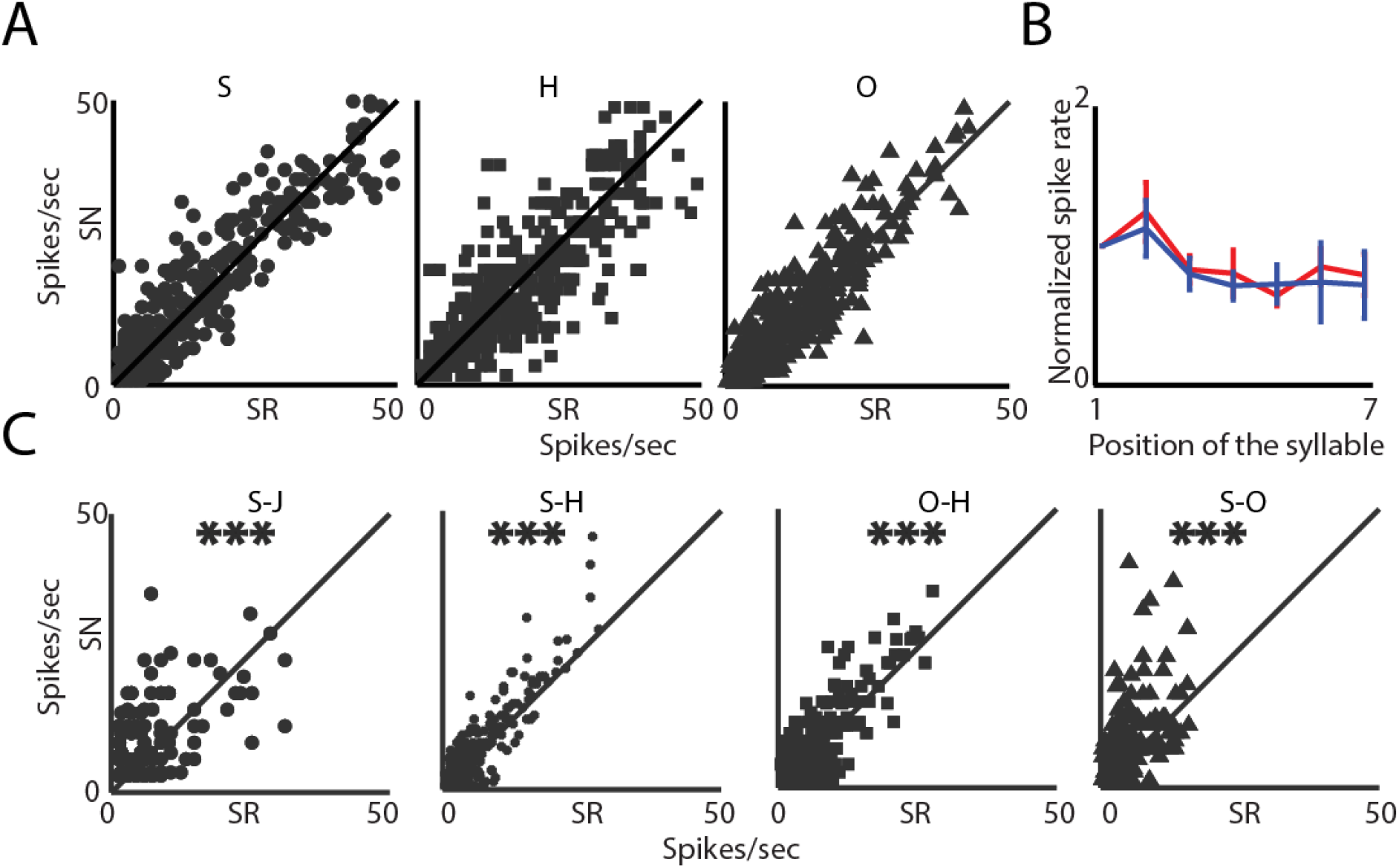
No alteration due to experience in mean response rate and time scales of adaptation. A) Scatter plots show comparison of mean response rates of all common first (identified by solid symbols, S: large circle, H: square, O: triangle) syllables types in SN and SR for AF-AFEx. B) Profile of changes in the normalized response strength for each position of the sequences SN (blue) and SR (red) over time with respect to the syllable in the starting position. C) Scatter plot of mean rate responses to first transition in SN (S-J, S-H, S-O and O-H) based on response to the second component, excluding the first transition.

**Figure S5.**
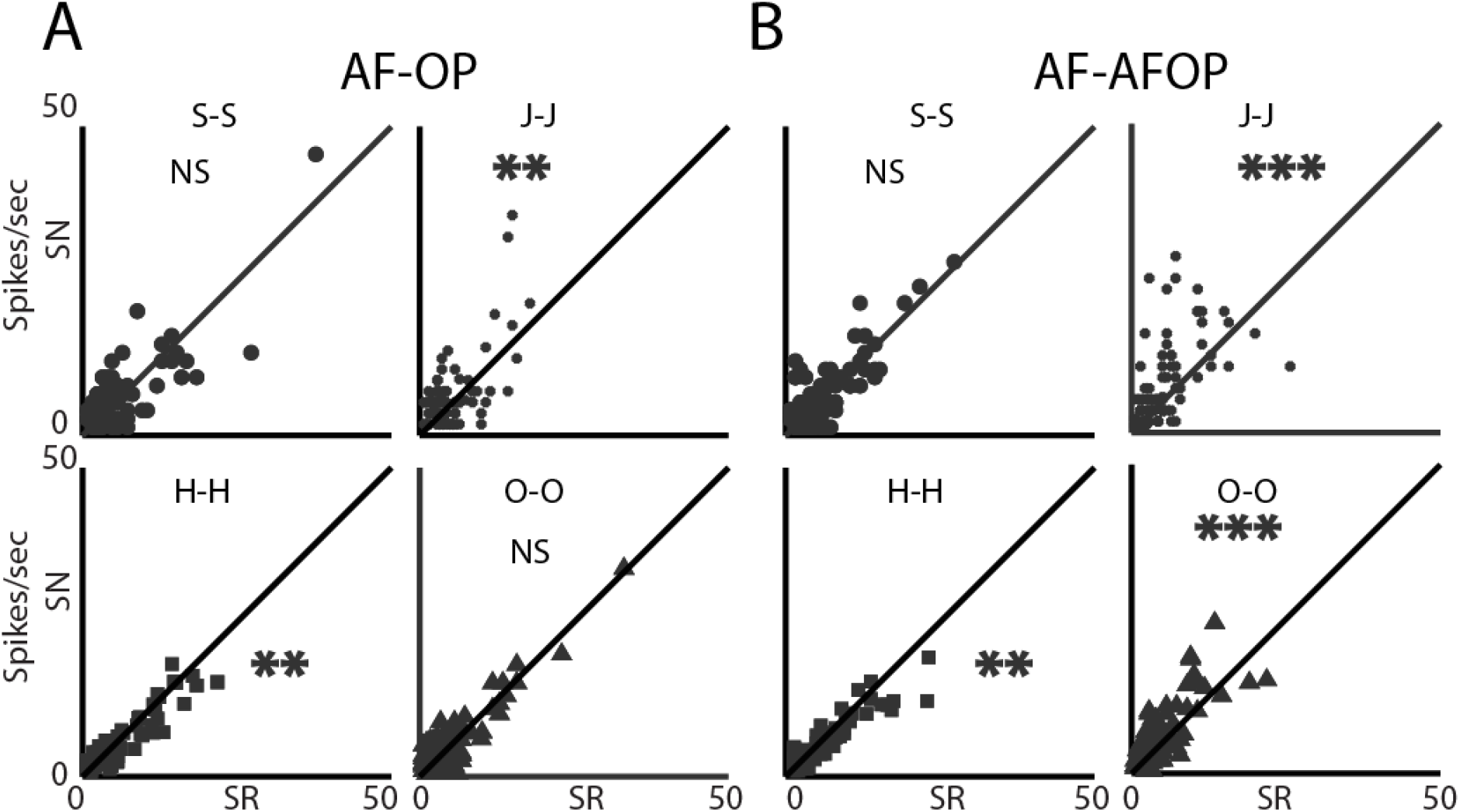
Effect of plasticity due to photo inhibition of SOM not reflected at disyllabic level. AB) Scatter plots comparing mean response rates of common syllables (identified by solid symbols, S: large circle, J: small circle, H: square, O: triangle) in SN and SR, excluding occurrence in the first position for the AF-OP and AF-AFOP groups. The arrangement is same as in Fig.3B.

